# UNCONVENTIONAL INTERLEUKIN-1 SIGNALING IN CARDIAC DYSFUNCTION

**DOI:** 10.1101/2025.03.09.642166

**Authors:** Stefano Toldo, Pratyush Narayan, Eleonora Mezzaroma, Alessandra Ghigo, Federico Damilano, Carlo Marchetti, Adolfo Gabriele Mauro, Emilio Hirsch, Benjamin Van Tassell, Antonio Abbate

## Abstract

Interleukin-1β (IL-1β) is an apical pro-inflammatory cytokine that has also been shown to negatively modulate cardiac contractility. Whether IL-1β effects on systemic inflammation and cardiac function are intertwined and associated with each other, or whether they are independent of each other, is unknown. An unconventional signaling of the IL-1 receptor type I through the phosphoinositide-3 kinase *γ* (PI3K*γ*), at least in part independent of the proinflammatory signaling, has been characterized in inflammation and cancer. We hypothesized that IL-1β would increase the expression of PI3K p110*γ* in cardiomyocytes, which in turn results in selective induction of p87 co-signaling and cardiac dysfunction through a scaffolding function on phosphodiesterase 3B (PDE3B). Using genetically modified mice, we show that a kinase-independent PI3K p110*γ* mechanism mediates IL-1-induced cardiac dysfunction. This may have compelling implications for the understanding and treatment of heart failure with reduced ejection fraction.

## TEXT

Heart failure with reduced ejection fraction (HFrEF) is associated with a systemic inflammatory response^1^. Interleukin-1β (IL-1β) is an apical pro-inflammatory cytokine that has also been shown to negatively modulate cardiac contractility^1^. Whether IL-1β effects on systemic inflammation and cardiac function are intertwined and associated with each other, or whether they are independent of each other, is unknown. The systemic effects of IL-1β are mediated through the regulation of nuclear factor *k*B (NF-*k*B) activation, which modulates the expression of secondary mediators, including Interleukin-6 (IL-6). Inhibition of IL-1β results in reduced systemic inflammation and reduced HF-related events.^1^ An unconventional signaling of the IL-1 receptor type I through the phosphoinositide-3 kinase *γ* (PI3K*γ*), at least in part independent of NF-*k*B and IL-6, has been characterized in inflammation and cancer.^2^ PI3K*γ* is a pro-inflammatory kinase involved in leukocyte function. Its expression is also increased in cardiomyocytes in HFrEF, and it has been shown to modulate cardiac contractility through a scaffolding, kinase-independent mechanism.^3-4^ Whether the cardiodepressant effects of IL-1β signaling are mediated through an increased expression of PI3K*γ* remains unknown. We hypothesized that IL-1β would increase the expression of PI3K *p110γ* in cardiomyocytes, which in turn results in selective induction of *p87* co-signaling and cardiac dysfunction through a scaffolding function on phosphodiesterase 3B (PDE3B) (**Figure, panel A**).

**Figure. 1.**
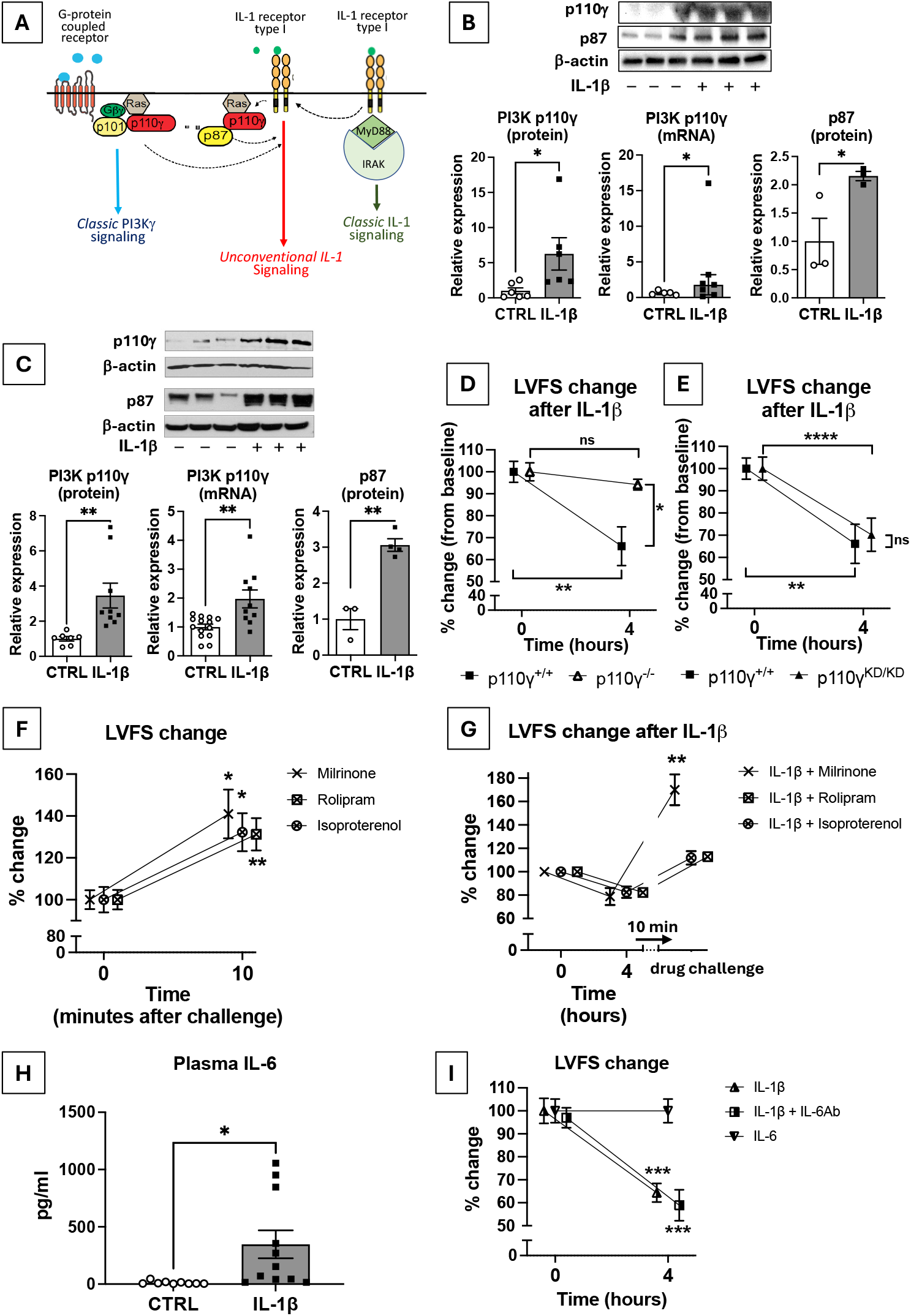
(A) Graphic representation of an unconventional IL-1 signaling converging on PI3K*γ* in the heart. (B) Adult primary cardiomyocytes were incubated with PBS (control) or recombinant murine (rm) IL-1β (50 ng/ml) for 4 hours; p110*γ* protein (N=6/group), p110*γ* mRNA (N=5-7/group), and p87 protein (N=3/group) expression were measured, *p<0.05. (C) Adult mice were treated with PBS (control) or rmIL-1β (3 *μ*g/ml) for 4 hours;p110*γ* protein (N=7-9/group), p110*γ* mRNA (N=10-14/group), and p87 protein (N=3-4/group) expression were measured, **p<0.01. (D) Adult PI3K p110*γ*^+/+^ (N=7) and p110*γ*^-/-^ (N=4) mice were treated with rmIL-1β (3 *μ*g/ml) for 4 hours; the cardiac function was measured at baseline and after 4 hours. *p<0.05; **p<0.01. (E) Adult PI3K p110*γ*^+/+^ (N=7) and p110*γ*^KD/KD^ (N=13) mice were treated with rmIL-1β (3 *μ*g/ml) for 4 hours; the cardiac function was measured at baseline and after 4 hours. **p<0.01; ****p<0.0001. (F) Adult wild type mice (N=5-7/group) were treated with Milrinone (PDE3 inhibitor, 10 mg/kg, N=5), rolipram (PDE4 inhibitor, 10 mg/kg, N=7), or isoproterenol (300 ng/kg, N=6); the cardiac function was measured at baseline and after 10 minutes. *p<0.05 and **p<0.01 vs baseline. (G) Adult wild type mice received rmIL-1β (3 *μ*g/ml) for 4 hours before rescue with Milrinone (PDE3 inhibitor, 10 mg/kg, N=4), rolipram (PDE4 inhibitor, 10 mg/kg, N=5), or isoproterenol (300 ng/kg, N=5); the cardiac function was measured at baseline, 4 hours after rmIL-1β injection and 10 minutes after the challenge with the PDE inhibitors or isoproterenol. **p<0.01 vs rolipram and isoproterenol treatment. (H) Adult wild-type mice were treated with PBS (control, N=9) or rmIL-1β (3 *μ*g/ml, N=11) for 4 hours to measure IL-6 plasma concentration, *p<0.05. (I) Adult wild-type mice were treated with rmIL-1β (3 *μ*g/ml, N=5), rmIL-1β and a blocking antibody against mouse IL-6 (5 mg/kg 12 hours before IL-1β, N=7) or rmIL-6; the cardiac function was measured at baseline and after 4 hours from rmIL-1β or rmIL-6 injection. ***p<0.001; vs baseline. *Abbreviations: Gβγ, G-protein coupled receptor βγ regulatory protein; IL-1, interleukin 1; IRAK, Interleukin-1 associated Kinase; LVFS, left ventricular fractional shortening; MyD88, myeloid differentiation factor 88; PI3Kγ, phosphoinositide-3 kinase γ; p110γ, catalytic subunit of PI3Kγ; p87, regulatory subunit of PI3Kγ; p101, regulatory subunit of PI3Kγ; p110γ*^*KD/KD*^, *p110γ kinase dead*.

To test this hypothesis, we isolated primary cardiomyocytes from adult mouse hearts using a Langendorff perfusion digestion with collagenase-2 and protease-IV. We determined that IL-1β (50 ng/ml for 4 hours) had no effects on viability assessed with trypan blue (data not shown) while it significantly induced a 6.2 and a 4.4-fold increase in *p110γ* protein and mRNA, respectively. Additionally, the *in vitro p87* protein expression was increased 2-fold (**Figure, panel B**). Using an established model of IL-1-induced cardiac dysfunction,^5^ we show that IL-1β (3 *μ*g/kg intraperitoneal) led to a significant reduction in left ventricular ejection fraction (LVEF) measured by M-mode echocardiography in the mouse after 4 hours. This was associated with an increase in PI3K *p110γ* protein and mRNA expression, 3.5-fold and 2-fold, respectively (**Figure, panel C)**, while *p87* protein expression increased by 3-fold. On the contrary, genetic deletion of PI3K *p110γ* (Pik3cg^tm1Dwu^) prevented dysfunction in mice treated with IL-1 (**Figure, panel D**). Furthermore, we showed that IL-1-induced cardiac dysfunction was maintained in a genetically modified mouse model that expressed a structurally intact PI3K *p110γ*, but catalytically inactive kinase - *p110γ* kinase-dead (KD) mice (PI3K*γ*^KD/KD^) - supporting that the scaffolding function is sufficient for IL-1-induced cardiac dysfunction (**Figure, panel E**).

PI3K p110*γ* is known for serving as a scaffold for PDE3B, a known regulator of cardiac contractility.^3^ We thus tested a dose-response in healthy mice for milrinone and rolipram, pharmacologic inhibitors with preferential PDE3 and PDE4 inhibition, respectively. A 10 mg/kg dose of milrinone and rolipram, increased fractional shortening (FS) by +40±10% and +31±8%, respectively. We also used isoproterenol, a β-adrenergic receptor agonist, as a control and chose a dose (0.3 µg/kg) associated with a similar response in FS (+32±8%) (**Figure, panel F**). We then treated mice with IL-1β and after 4 hours attempted to rescue them with milrinone, rolipram, or isoproterenol. Only treatment with milrinone entirely reversed IL-1-induced dysfunction (**Figure, panel G**).

In a separate set of experiments, we tested whether blocking IL-6 would also be sufficient in preventing IL-1-induced cardiac dysfunction. Using the same model of IL-1-induced cardiac dysfunction, we found that while IL-1β induced a significant increase in serum IL-6 levels (**Figure, panel H**), pre-treatment with previously validated IL-6 neutralizing antibody (5 mg/kg administered intraperitoneal 12 hours before IL-1 β) did not prevent cardiac dysfunction, thus supporting that IL-1-induced cardiac dysfunction is not mediated by circulating IL-6 (**Figure, panel I**).

Determining the mechanism by which IL-1β modulates cardiac function independent of systemic inflammation has a potential impact on drug development. IL-1β inhibition has shown to be beneficial in patients with or at risk for HFrEF, however this was associated with a small, yet significant, increase in fatal infections ^1^. Defining how PI3K*γ* participates in IL-1 signaling in cardiomyocytes and contributes to cardiac dysfunction may open the way to more targeted therapies to prevent and treat HFrEF. The distinction between kinase-dependent and -independent functions of PI3K *p110γ* in IL-1-induced cardiac dysfunction provides an additional opportunity to differentiate pro-inflammatory and cardiodepressant mechanisms in HFrEF. The possibility that the deleterious effects of IL-1β on cardiac function are independent of IL-6 raises questions regarding whether or not IL-6-targeted strategies would prove beneficial in HFrEF. Of note, with its complex signaling, IL-6 has been described as a double-edged sword in HFrEF.

In conclusion, we show that a kinase-independent PI3K p110*γ* mechanism mediates IL-1-induced cardiac dysfunction. This may have compelling implications for the understanding and treatment of HFrEF.

## Abbreviations

IL-1β: Interleukin-1β
NF-*k*B: nuclear factor *k*B
IL-6: Interleukin-6
PI3K*γ*: phosphoinositide-3 kinase *γ*
p110*γ*: catalitic subunit of PI3K*γ*
KD: Kinase dead
p87: regulatory subunit of PI3K*γ*
HFrEF: heart failure with reduced ejection fraction
PDE3: phosphodiesterase PDE3
PDE4: phosphodiesterase PDE4

## Conflict of Interest Disclosures

Stefano Toldo: CardiolRx, Monte Rosa Therapeutics (consulting).

Alessandra Ghigo and Emilio Hirsch: Kither Biotech Srl (co-founders and shareholders). Federico Damilano is currently employed at Pfizer Inc. (Cambridge, MA, USA), but his contribution to this report was conducted during his affiliation to UniTo.

Antonio Abbate: CardiolRx, Kiniksa Pharmaceuticals (consulting).

## Acknowledgements

We thank Dr. Howard Rockman for providing Pik3cg^tm1Dwu^ mice (PI3K *p110γ*^-/-^).

## Funding Sources

The study was supported by a NHLBI grant to Stefano Toldo and Antonio Abbate (R01HL150115).

## Data availability

Upon request by e-mail, Stefano Toldo, PhD within 24 months of study completion, will provide access to individual experimental data.

